# Flocked swab might be one main reason causing the high false-negative rate in COVID-19 screening----the advantages of a novel silicone swab

**DOI:** 10.1101/2020.03.29.014415

**Authors:** Jianye Zhou, Zhongtian Bai, Xiaoping Liu, Yaqiong Guo, Nan Jiang, Xiaodong Li, Xiaohui Zhang, Zhiqiang Li, Yonghong Li, Zhongren Ma, Jin Zhao

**Affiliations:** Biomedical Research Center, Key Laboratory of Oral Diseases of Gansu Province, Key Laboratory of Stomatology of State Ethnic Affairs Commission, Northwest Minzu University, Lanzhou, Gansu Province, China; The second Department of General Surgery, the First Hospital of Lanzhou University, Lanzhou, Gansu Province, China; Department of Respiratory Medicine, Second People’s Hospital of Gansu Province & Northwest Minzu University Affiliated Hospital, Lanzhou, Gansu Province, China; Department of Applied Soil Biochemistry, Institute of Applied Ecology, Chinese Academy of Sciences, Shenyang Province, China; Institute of Chinese Materia Medica, Gansu Provincial Hospital of Traditional Chinese Medicine, Lanzhou, Gansu Province, China; NHC Key Laboratory of Diagnosis and Therapy of Gastrointestinal Tumor, Gansu Provincial Hospital, Lanzhou, Gansu Province, China

**Keywords:** COVID-19, Specimen collection, RNA extraction, swab

## Abstract

RNA testing using RT-PCR can provide direct evidence for diagnoses of COVID-19 which has brought unexpected disasters and changes to our human society. However, the absorption of cotton swab for RNA lysates may lead to a low concentration of detectable RNA, which might be one of the main reasons for the unstable positive detecting rate. We designed and manufactured a kind of silicone swab with concave-convex structure, and further compared the effects of silicone and cotton swab on RNA extraction. Principal component analysis and Paired Wilcoxcon test suggested that a higher RNA concentration and A260/A280 would be obtained using silicone swab. The results indicated that our silicone swab had a more excellent ability to sample than the cotton swab, characterized by the higher quantity and quality of extracted RNA. Thus, we advised that the current cotton swabs need to be improved urgently in COVID-19 diagnoses and the process of “sample collection” and “sample pre-processing” must be standardized and emphasized.

**Highlights:** The current cotton swabs need to be improved urgently in COVID-19 screening.

## INTRODUCTION

Coronavirus Disease 2019 (COVID-19) has become the number one public enemy in the world now, bringing unexpected disasters and changes to our human society. Viral RNA testing using RT-PCR was recommended to provide direct evidence in diagnoses of COVID-19 [1]. However, the positive rate is frustrating (about 30%-60%), combined with a high false negative rate [2, 3], which create huge obstacles for the control and prevention of COVID-19. During the screening and detection for suspected cases, confirmed cases and close contacts, the preferred sequence of sample collection is nasopharyngeal swab, oropharyngeal swab and sputum [4]. Flocked swab is the standard material collecting samples for the first two [5]. The flocked swab stays in nasopharyngeal or oropharyngeal for 15-30 seconds, rotates gently 3-5 times, lyses and then extracts RNA [4]. However, importantly, the absorption of flocked swab for RNA lysates may lead to a low concentration of detectable RNA, which might be one of the main reasons for the low positive detecting rate. In addition, prolonged irritating with a throat swab causes subjects vomiting or coughing in practice, which would increase the risk of transmission and infection. However, to our knowledge, no studies have comparatively quantified the testing defect caused by using flocked swab, and no optimized product has been designed yet. Therefore, designing a novel swab owning abilities of both the stronger adhesion and more efficient releasing for samples might contribute to improving virus detection rate.

## RESULTS and DISCUSSION

In the process of coping with the major public health emergencies, such as COVID-19, increasing the real positive rate and reducing the false negative rate is crucial. The nuclear acid detection of pharyngeal mucous collected by flocked swab is the most convenient and common method. In order to reduce the false negative rate during clinical testing, updating novel detection techniques is always attracted the most attention of researchers, but evaluating the quantity and quality of nuclear acid has always been neglected. Several defects of flocked swabs had been found in our clinic application, which might be one of the main reasons for the high false negative rate. First of all, the flocked swab owns a strong absorption of mucous, but meanwhile, the soaked cotton wool becomes fluffy and samples could be deeply adsorbed. Thus, it is difficult to elute samples into lysis solution. Secondly, the flocked swab needs to be vibrated using steel beads for collecting viral cells as many as possible in laboratory, but it is too cumbersome to operate in clinical testing. Thus, thirdly, the swab needs to be stretched into the oropharynx and wiped repeatedly to collect more. However, the ligneous stick and the hard flocking surface of the swab would cause resistance of the subjects due to pain or discomfort. Hence, the collecting result may be not satisfactory, which to a certain extent, results in missing detection.

Therefore, we produced a novel swab made of elastic silicone gel, and subtly designed a concave-convex structure for improving the adhesion and increasing the adhesion surface (Fig S1). The elastic silicone gel is obviously softer and more comfortable than the flocked swab. In addition, the mucus mixture will be collected more efficiently on the swab surface and also be eluted more easily using our silicone swab than the cotton swab. Based on the examined indexes, including the concentration (i.e., quantity) as well as A260/A280 (i.e., quality), of extracted RNA, PCA showed that the individuals could be extremely well separated by the silicone and cotton groups (*P* < 0.001, Fig 1A). In particular, the PC1 explaining 99.7% of the variations in data showed an extremely significant difference between the cotton and silicone groups (*P* < 0.001, inserted in Fig 1A). Specifically, the concentration of RNA was significantly higher from the silicone group than the cotton group based on paired comparison (*P* < 0.001, Fig 1B), separately accounting for 9.4 and 6.4 ng/μl for the silicone and cotton group. In addition, the value of 260/280 which represents the purity of nucleic acids was also significantly higher in the silicone group (*P* = 0.01, Fig 1C). These results indicated that the elastic silicone gel significantly improved sampling efficiency.

**Figure 1:**
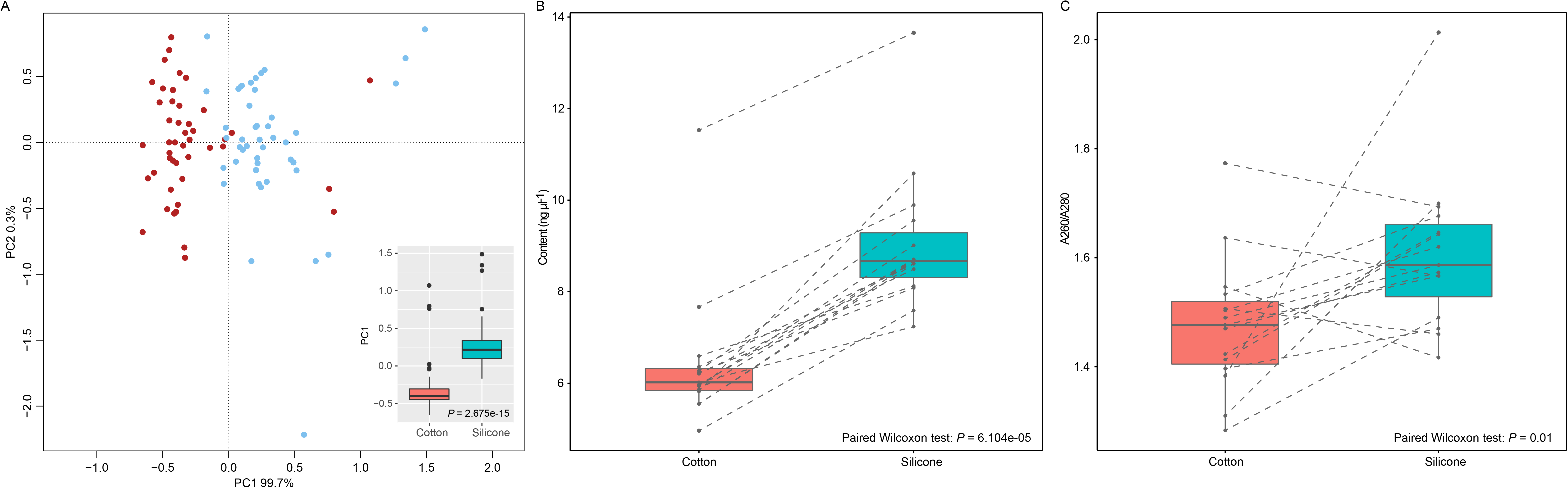
PCA analysis including content, A260/A280 and the comparing of content Comparisons of content and quantity and quality of extracted RNA. Blue and Red indicate the cotton and silicone groups, respectively. (A) Principal Components Analysis of data, including RNA content and A260/A280, was performed across 15 individuals (each individual in triplicate). The comparison of the values of PC1 was calculated based on Wilcoxon test and inserted as a boxplot. The comparison of RNA concentration (B) and A260/A280 (C) between the cotton and silicone groups based on paired Wilcoxon test. The dotted line indicates the data pair of cotton and silicone for each individual.

Last but not least, the infection of COVID-19 for medical staff occurs in all aspects of prevention and treatment. For viral RNA testing, after sampled, the swab applicator was cut and placed into a tube containing RNA preservation solution. Then the sealed sample tube was delivered to the clinical lab for detection. Although RNA preservation solution can lyse the viral shell and make it inactive, the residuary flocked swab increased the infection risk of the examiner. This defect could be avoided by using the elastic silicone swab. The samples collected by the elastic silicone swab would be almost fully eluted into the tube, and the swab would be discarded as medical waste. Therefore, the detectors would not contact with the polluted swabs, minimizing the infection risk of COVID-19.

Overall, we advise clinicians to pay more attention to the process of mucus collection, which might significantly improve the early detection rate of COVID-19. It’s worth mentioning that during the critical period confronting COVID-19 globally, we decided to announce the findings in time, even though the results were not verified by a big data. Even so, our data suggested that the current swabs need to be improved urgently. In the following research, we will optimize the elastic silicone gel and conduct comprehensive index detection so as to reduce the high false negative rate due to the defect of the sampling swab.

In addtion, there are some advises about the COVID-19 screening in the present study:

1. The sample collection and sample pre-processing before the RNA extraction should be standardized and emphasized so as to eliminate the false-negative as much as possible.
2. In the process of sample collecting, whether the sampling area is nasopharyngeal or palatopharyngeal, the following three important factors should be paid attention: sufficient contact with sampling area; a certain strength of sampling and a sufficient sampling duration. The swab which owning excessive interstices like flocking swab should be avoided. To ensure that more samples are obtained during extraction, the swab owing higher rates of adhesion and elution is recommended.
3. In the process of sample pre-processing, the swab head should be adequately pressed on the inner wall of the centrifuge tube and wash swings repeatedly so that the sample can be fully eluted into the RNA lysate. Befor using RNA extraction Kit, a step of centrifugation with 13,000rpm at 10min must be stipulated for dealing with all eluant and collecting the sediment, rather than sucking a portion of the eluent without centrifugation current widely used. During this process, the centrifuge should be ask for putting into biosafety cabinet to avoid aerosol infection.

## Methods

### Sample collection and RNA extraction

We designed and manufactured a kind of silicone swab with concave-convex structure and compared its effects on RNA extraction with cotton swab provided by Department of Respiratory Medicine, Second People's Hospital of Gansu Province & Northwest Minzu University Affiliated Hospital, Lanzhou, Gansu Province, China. 15 volunteers were from the Hospital too. They were recruited and accepted the pharyngeal examination. The tumors, lesions and hyperplasia in laryngeal part of pharynx were excluded. The Pharyngeal mucous were sampled by an experienced technician who were passed the specifically bio-safety training held by the management department of the affiliated hospital of northwest university for nationalities. The technician was required equipped with protective equipment according to the three levels of protection according to the third level biological safety standard, including work clothes, protective clothing, disposable working cap, medical protective mask (N95), surgical mask, goggles, mask, double gloves, waterproof boots and shoe covers. The oropharyngeal palatum was divided into the same two parts by visualization, one of which was sampled with flocked swab, and the other was sampled with silicone swab. The swab stayed in the sampling area for 15-30 seconds, and was gently rotated for 3-5 times. After the samples were obtained, both kinds of swabs were placed separately into two tubes containing lysis solution (Guangzhou BDS Biological Technology Co., Ltd). Then, the swab was repeatedly squeezed on the inner wall of the collection tube for 10 times and removed, so that the sample can be rinsed into the lysis solution as much as possible. Then screwed the tube and sealed it for laboratory testing. The RNA extraction process was referred to the instructions of the kit (RNeasy Mini kit, QIAGEN, Germany). The extracted RNA was detected by ultra-fine ultraviolet spectrophotometer (NanoDrop, Thermofisher, USA), three times for each sample. The protocol for this study was approved by the Ethics Committee of the Northwest Minzu University.

### Statistical methods

The comparison between the silicone group and the cotton group were analyzed by a principal component analysis (PCA) across the samples using the vegan package in R (version 3.6.2). Wilcoxon test was performed to compare PC1 between the two groups using the vegan package in R, as to PC1 with a nonnormal distribution. In addition, paired Wilcoxon tests were performed and visualized using ggpubr package in R to compare the content of RNA and the values of 260/280, both of which were nonnormal distributions, from the 15 individuals.

## Acknowledgments

We are extremely grateful to all the volunteers in the present study and appreciate the Prof. Qiuwei Pan who gave advice on the study design.

## Author Contributions

JY. Z designed this experiment and produced the silicone swab. ZT. B involved in the design and demonstrated the manuscript. XP. L carried on the clinical sampling, and YQ. G extracted and examined the RNA. N. J performed statistical analyses of the data and plotted the results. XD. L and XH. Z assisted the manufacture of silicone swab. YH. L guided the process of sampling and extracted the RNA. ZQ. L guided the shape of the silicone swab. ZR. M and J. Z guided this research and reviewed the manuscript.

## Declaration of Interests

The authors declare that the research was conducted in the absence of any commercial or financial relationships that could be construed as a potential conflict of interest.

## Funding

This study was supported financially by projects of the double first-class and special development guidance, Northwest Minzu University (11080306-2019) and National Natural Science Fund of China (81702326)

